# Estimating home range size under spatial constraints: a comparative approach using the semi-aquatic European mink (*Mustela lutreola*)

**DOI:** 10.64898/2026.04.13.718143

**Authors:** Rémi Bodinier, Stéphane Aulagnier, Yoann Bressan, Romain Beaubert, Christine Fournier-Chambrillon, Sébastien Devillard, Pascal Fournier

## Abstract

Accurate home range knowledge is essential for conserving species that are highly constrained by spatial features. The critically endangered European mink (*Mustela lutreola*) is a wetland specialist whose movements are constrained along rivers or in wetlands. In dendritic landscapes, conventional home range estimators such as Minimum Convex Polygons tend to include unsuitable areas in estimated home ranges. Using VHF telemetry data from 16 individual-years tracked in France between 1996–1999 and 2020–2022, we compared four methods: Kernel Density Estimator (KDE), an adaptative sphere-of-influence local convex hull (a-LoCoH), a newly developed Ecological Home Range method (EHR), and a Generalized Additive Model (GAM) approach integrating hydrographic covariates. Our objective is to determine which method best accounts for the European mink’s specialization in wetlands, considering the spatial distribution of locations. Evaluation with a wetland-specific metric showed KDE consistently overestimated range extent and included unsuitable areas, and a-LoCoH yielded mixed results, but these indicated that the method was not effective in excluding unused areas. It was EHR and GAM methods that aligned more closely with ecological constraints. We therefore recommend GAM because it matches our objective and has the capacity to integrate additional environmental variables. Using the GAM, male home ranges averaged 3,074 ha—26 times larger than female ranges (116 ha)—and were significantly larger in river than marsh landscapes. These are the largest ranges reported for the species. Large spatial requirements heighten vulnerability to road fatality and predation, both significant threats for remaining French populations. Our findings highlight the need for conservation strategies that integrate precise, spatial-constraint-based range estimates. The GAM method offers a robust, adaptable framework for managing European mink and other semi-aquatic species in complex landscapes.

## 1. Introduction

According to Burt (1943)’s definition, the home range is the area traversed by the individual during its normal activities of food gathering, mating, and caring for young, excluding occasional exploratory sallies. Home range mapping is therefore essential for conservation planning of endangered species. So, identifying home range characteristics helps informing management strategies, and guiding protection measures of both species and the areas in which they live (Fleming et al., 2015; Goldingay, 2015). The way in which an animal uses its surrounding environment for its daily activities may reflect continuous use of the entire space or more patchy use with areas of greatest use separated by areas which the animal regularly passes through without stopping. In all cases, the home range must correspond to the ecological requirements of the species. It is particularly true for populations that are restricted to dendritic systems and estimating the extent of home ranges presents unique challenges (Kenward, 2000; Ouellette and Cardille, 2011). In dendritic landscapes, like rivers for example, movement is typically associated with the branching structure of watercourses inside the wetlands, leading to spatially structured dispersal patterns (Sutherland et al., 2015). Yet, most traditional home range estimation methods do not consider the spatial constraints associated with such dendritic systems.

This is mostly relevant for highly declining species of high conservation concern such as the European mink *Mustela lutreola* (Maizeret et al., 2002). It is classified as critically endangered (CR) by the IUCN and is one of Europe’s most endangered carnivores (Maran et al., 2016). The species is highly sensitive to habitat loss, overexploitation, road kills and invasive species such as the alien American mink *Mustela vison* (Maran and Henttonen, 1995; Maran et al., 2009; Maran et al., 2017; Põdra and Gómez, 2018). Additional threats include predation by red foxes (*Vulpes vulpes*), domestic dogs, and raptors (Maran et al., 2009; Maran et al., 2017).

The European mink is strongly associated with watercourses and wetlands. It may explore entire river basins but rarely strays more than 150 metres from watercourses (Danilov and Tumanov, 1976), indicating that proximity to water strongly constrains the space used by the species. Within these broad areas, the European mink uses a variety of water-associated environments, including rivers, streams, ponds, canals and marshes, with some areas being used more extensively than others (Zabala and Zuberogoitia, 2003; Fournier et al., 2007, Palomares et al., 2017a). Accordingly, its home range is expected to be closely linked to the spatial distribution of watercourses and other wetland features, rather than to the overall extent of the surrounding landscape (Fournier et al., 2008; Palomares et al., 2017b). This strong spatial association with water has important implications for home range estimation. In the case of the European mink, inappropriate models may fail to capture the spatial constraints imposed by its dependence on water-associated environments, potentially leading to inaccurate estimates of home range size and shape, misidentification of key areas, or underestimation of connectivity needs. The acquisition of new data in France and the emergence of more effective home range estimation methods therefore make understanding its spatial requirements crucial for targeted conservation measures. Consequently, there is a growing need for methods that incorporate variables reflecting the species’ dependence on water and wetlands, while accounting for the spatial structure and connectivity of the systems it uses.

Based on VHF (Very High Frequency) telemetry data collected in two French conservation projects (first: from 1996 to 1999; second: from 2020 to 2022) deployed in riverbank and marsh landscapes, we evaluated four methods to describe European mink home range. Our main objective is to identify the best method for estimating home range size, taking into account both spatial constraints, i.e. the European mink’s high specialization in wetlands, and the nature of our data, i.e. daily locations. We will therefore seek the method that best matches the spatial definition of the wetland while allowing for the possibility that this home range may be estimated as continuous or patchy. We have decided to rule out the Minimum Convex Polygon (MCP –Mohr, 1947), the most commonly used home range estimator method, because this method tends to overestimate home range size by including unused or inaccessible areas, especially in linear landscapes (Slaght et al., 2013; Royle et al., 2013). We therefore opted to compare two reference known control methods to two hydrographic covariates-based methods. As reference we chose the kernel density estimator method (KDE –Worton, 1989) because of its widespread use in home range modelling, and the adaptative sphere-of-influence Local Convex Hull (a-LoCoH –Getz et al., 2007) which is a method that is increasingly highlighted in the literature. The two methods we developed are, for the first one named the Ecological Home Range (EHR), based on locations distribution along watercourses, and the second named GAM, coming from the species distribution modelling framework. Each method was also used to estimate the core area; core area being defined as area of greater use within the boundaries of the home range.

Our aim was thus to investigate the strengths and limitations of these methods in representing wetland-constrained home range of a highly threatened semi-aquatic species. Although it is a highly adaptable method, the Kernel method is likely to be the least effective for modelling the home range of the European mink,as its known poor ability in retrieving constrained home-range (Blundell et al., 2001). The a-LoCoH method should be better at modelling the home range of the European mink because it has been shown that this method is more effective at modelling holes in the home range in unused areas (Lichti and Swihart, 2011; Chirima and Owen-Smith, 2015). Ultimately, the two methods we have developed are likely to be the best methods for estimating home range, as they take into account the ecological constraints specific to the species. Indeed, these methods are based on hydrological variables, which are the primary factor in determining the home range of the European mink. Once the best objective-dependent method has been chosen, we investigated the pattern of variability of home range size and core area according to sex, season and landscape type to provide key information on the amount of space used by the European mink and, ultimately, recommendations for its conservation, benefiting for other wetland-dependent mammals.

## 2. Methods

### 2.1. Study area

We conducted our study in three study sites (**Fig. 1**).

**Fig. 1:**
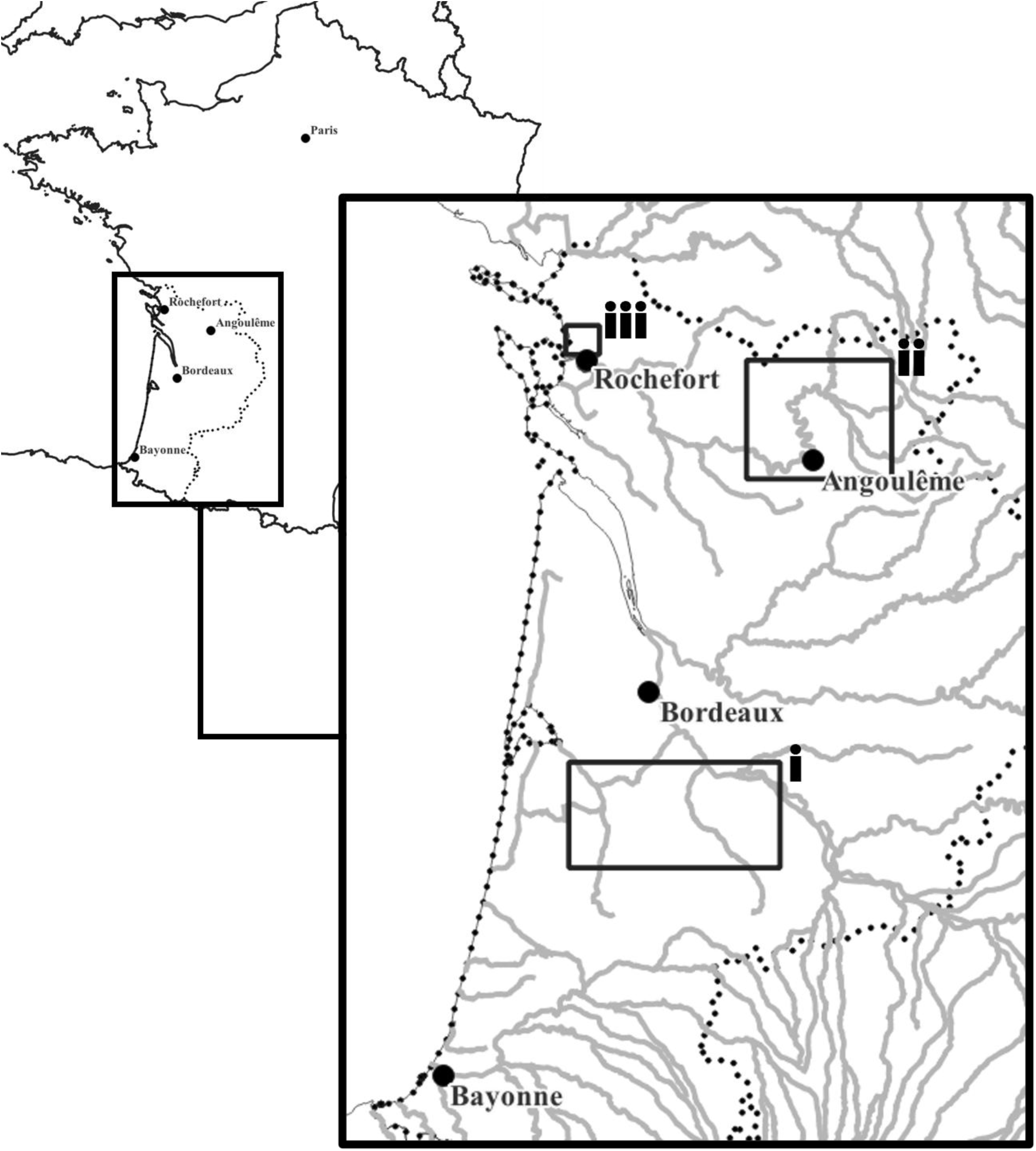
Study area. i) Landes de Gascogne region (LG), ii) Charente River catchment upstream from Angouleme (CA), iii) Rochefort marsh (RM). Grey lines represent watercourses, and black dotted lines represent the boundaries of French departments where European was suspected to occur at the beginning of the 21^st^ century.

First, the Landes de Gascogne region (LG; 44°22’N, 0°34’W, 1996-1999) in southwestern France covers over 10 000 km², mainly occupied by highly productive *Pinus pinaster* plantations. River valleys unsuitable for intensive forestry host unexploited deciduous forests and swamps. Fieldwork focused on the Eyre and Ciron river valleys. Second, the Charente River catchment upstream from Angouleme (CA; 45°38’N, 0°20’E, 2020-2022) spans 3500 km², one third covered by cultivated lands. Natural habitats are mostly wet grasslands and deciduous forests. The landscape of LG and CA are considered ‘river’ type. Third, the Rochefort Marsh (RM; 46°1’N, 0°56’W, 2020-2022) is a backshore marsh of tidal mudflats and brackish hygrophilous meadows, crossed by a dense network of freshwater ditches. This area is considered a ‘marsh’ landscape type.

### 2.2. Captures and sampling

Captures and sampling procedures were similar between the two periods in the three study sites. Fournier et al. (2007; 2008) published detailed procedures for the LG study area. For both CA and RM study areas, we organized trapping campaigns from mid-September to the end of March, avoiding the breeding season. We set 40 to 50 non-commercial live traps (75 x 16 x 16 cm) baited with sardines along 5 to 10 km of rivers and streams or in 10 to 15 km² of marshes for at least 10 consecutive nights. Trapping refinement consisted of plastic protection against rain, hay for resting and a piece of 150g trout for energy boost. We captured European mink from 1996 to 1999 in LG, and from 2020 to 2022 in CA and RM. A total of 14,731 trap-nights in LG, 4498 in CA, and 4,793 in RM resulted in the capture of respectively 11 (4 ♀, 7 ♂), 5 (3 ♀, 2 ♂) and 7 (1 ♀, 6 ♂) individuals, all considered to be adults according to capture date and reproductive status, except for one male in CA, which was at least 6 to 8 months old and likely a subadult. Only mink in apparently good health received a transmitter, the other ones were released a few hours after the capture, after hair sampling and microchip tagging. We defined health as good or not with a threshold weight, taking into account natural seasonal variations (500 g for females and 800 g for males in autumn, 520 g for females and 875 g for males in winter).

We carried out further procedures in a dedicated room located near the capture site. Individuals were immobilized with 150 μg/kg of medetomidine (Domitor®, Vetoquinol, France) combined with 7.5 mg/kg of ketamine (Ketamine 1000®, Virbac, France) intramuscularly, antagonized by 750 μg/kg atipamezole (Antisedan®, Vetoquinol, France). They were placed under oxygen masks and on heat mats, were rehydrated subcutaneously with 4 ml/kg of a saline solution (Ringer Lactate Lavoisier®, Laboratoires Chaix et du Marais, France) and received 2 mg/kg Tramadol analgesic (Tralieve®, Dechra veterinary Products SAS, France) and 0.08 ml/kg of antibiotic (Shotapen®, Virbac, France). We used implantable transmitters provided by Telonics (Mesa, Arizona 85204-6699, USA) and sterilized with Ethylene Oxide: 16 g IMP-130 high and medium power range, with an expected operational life of 4.2 and 7.1 months respectively. Other procedures for fitted animals included a detailed clinical examination, hair sampling and microchip tagging. We released animals on their capture site about 24 hours after capture. French Ministry responsible for the Environment (Ministerial Decree of 19 February 2018) licensed captures and animal manipulations. Ethics committee COMETHEA CE084 and French Ministry responsible for Research (APAFIS file#15599-2018062011206645 v2) approved and authorized related procedures.

### 2.3. Radiotracking

We radio-tracked individuals from March 1996 to August 1999 in LG, and from February 2020 to May 2022 in CA and RM. In LG, we tracked two out of the 11 individuals captured during two different seasons and so we monitored them over two years (i.e. leading to two individual-years in the data). In CA, we tracked among four out of the five individuals trapped, and three during two different seasons. In RM, we monitored four out of the seven individuals trapped. On the whole, radiotracking resulted in monitoring 24 individual-years. We used TRX 1000S receivers (Wildlife Materials, Carbondale, Illinois 62901, USA) either connected to a 5- or 8-element yagi antenna mounted on a vehicle or to a 3-element yagi hand-held antenna, or a H antenna (RA-23 Telonics model). We tried to locate each animal daily, by triangulation from a vehicle and then, except if field conditions do not allow it or when the animal was found active, using the hand-held antenna by foot to know the precise location of the resting site. All the location data collected during daily monitoring were incorporated to estimate the individuals’ home ranges, in order to adhere as closely as possible to Burt’s definition. Indeed, amongst the location data used, some were recorded whilst the animal was active (when the signal was unstable –activity may in such cases involve movement or foraging), and a male and a female were found in the same roost during the breeding season (Bodinier et al., comm. pers.) and some females were tracked whilst rearing their young. Tracking lasted as long as the battery operated or as long as we found the individual alive.

### 2.4. Home range estimation

We estimated annual home range size following the four approaches below. All daily locations were used for each home range estimation, as other telemetry studies used the same type of data to estimate the home range of other semi-aquatic mammals (Leite et al., 2016; Esnaola et al., 2018; Halbrook and Petach, 2018).

First, we calculated fixed 95% Kernel Density Estimate (KDE) home range and 50% as core areas, using the adehabitatHR software package (Version 0.4.22, Calenge, 2015) in R software (Version 4.4.1, 2024-06-14, R Core Team 2016). Kernel estimation produces a smooth empirical Probability Density Function (PDF) based on individual locations.

Parameter *h*, the smoothing parameter, governs the amount of smoothing applied. We fixed this parameter for each individual as the mean Euclidean distance between each location and its nearest neighbour.

Second, we modelled the 95% adaptative Local Convex Hull (a-LoCoH) home range and the 50% one as core areas using also the adehabitatHR software package. This method uses all points within a variable circle around a root point such that the sum of the distances between these points and the root point is less than or equal to **a**. This method adjusts the radius of the circle that circumscribes each local convex hull, such that smaller convex hulls arise in high use areas. Following Getz et al. (2007), we fixed **a** as the maximum distance between any two points for each individual.

Third, we proposed a new method incorporating both location data and biologically relevant landscape features: specifically, the distance to the nearest watercourse, referred to as the Ecological Home Range (EHR). For each individual-year, we measured the distance from each location to the nearest watercourse, using the BD TOPO® hydrography dataset (version 3.3, Institut National de l’Information Géographique et Forestière). We delineated the watercourses network by cutting it perpendicularly to the most upstream and downstream relocations. We then created a buffer around all watercourses, using the maximum remaining distance after excluding the longest 5% of distances, assuming farther locations reflected exploratory behaviour and thus were not part of the home range. To define core areas, we constructed the median axis of the EHR and projected locations onto it. For each projected point, we measured its distance from a reference point (the most downstream location for each individual-year). We then estimated the probability density function of these distances with the kernel method, identifying the axis segments where probability equalled 50% as core areas. For each individual-year, we derived the smoothing parameter from that of the KDE-based home-range, then varied by ±10 m increments across five lower and five higher values. For each, we counted the number of core areas, the number of locations per core area, and the total number of locations. We then constructed the index *I:*

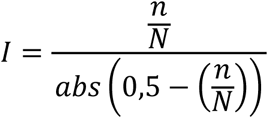

where *n* = minimal number of locations in core areas, and *N* = total number of locations.

High *I* value indicate both that the proportion of locations in core areas is close to 50%, and that each core area is supported by sufficient locations. We selected the smoothing parameter maximizing *I*.

Fourth, we applied a Generalized Additive Model (GAM) for each individual-year to explain distribution in presence/pseudo-absence data. We defined presence data as the individual-year locations. We created a square interest area (SIA) based on 1 km buffer around the northern, eastern, southern and western locations. We then randomly defined pseudo-absence points within this SIA. Since a large number of pseudo absence data are recommended in species distribution models (Stokland et al.,2011; Barbet-Massin et al., 2012), we created enough pseudo-absence points so that true locations represented 5% of the dataset (i.e. pseudo-absence = 95%). Following Stokland *et al*. (2011), the prevalence used to determine the absence of the species must match the ratio of presence points. We then defined the individual-year home range as the area modelled with a probability of presence > 0.05. The covariates in the GAM model included coordinates (in the form of a thin-plate spline) to account for the spatial distribution of individual locations. Furthermore, we included only hydrological covariates in the model, as access to water and wetlands is the main factor constraining the European mink’s use of space. These covariates are the distance from the nearest watercourse, the probability of being within a wetland, and the distance from the wetland’s median axis (**Table 1**). It should be noted that this last covariate could only be measured in riverine landscape (LG and CA) because, in marshy landscape, the globular shape of the wetland prevented the definition of such an axis. We constructed GAM-based core areas the same method as for the EHR: (i) modelling the median axis of the GAM-based home range, (ii) projecting points onto it and measuring distances from reference point, and (iii) calculating a 50% distance density estimate with the appropriate smoothing parameter.

**Table 1:**
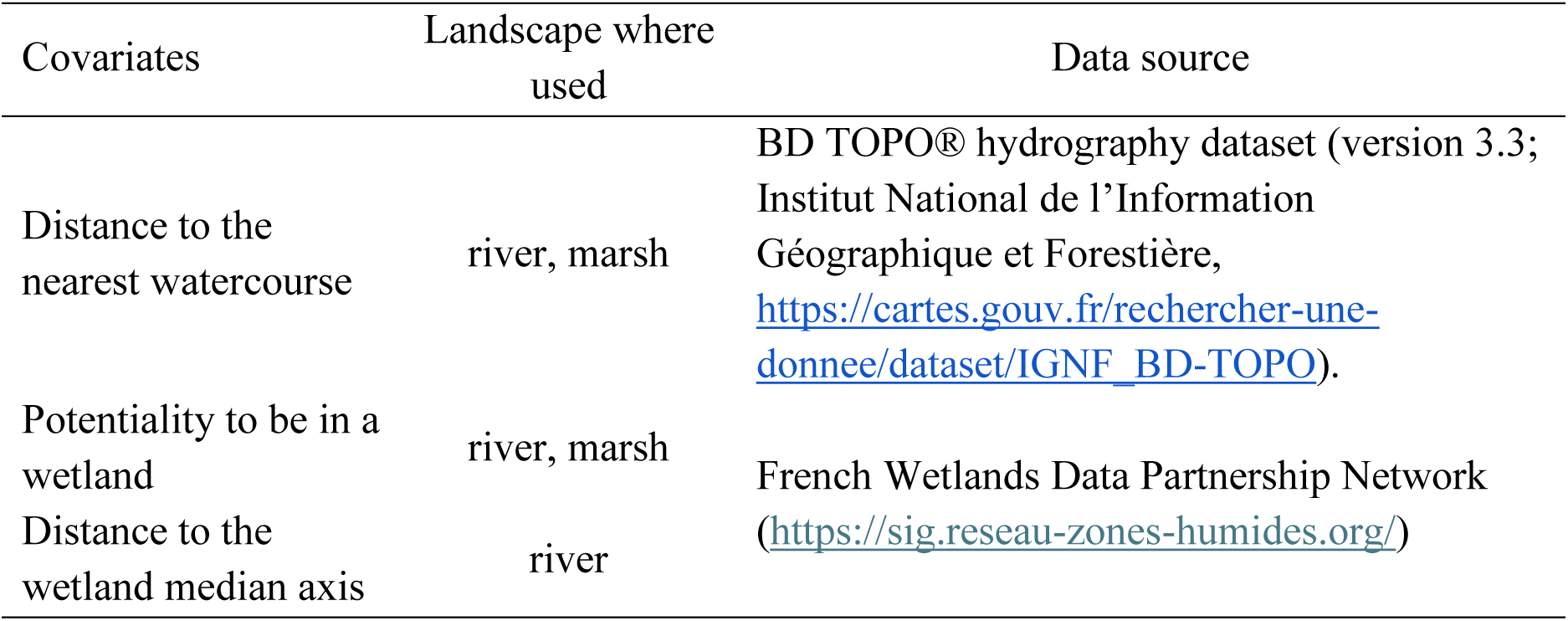
Hydrological covariates used in the GAM model to assess the European mink range and data source.

### 2.5. Reliability of home range estimators

We defined as a metric to compare the methods, reflecting strongly European mink specialisation to wetlands, the proportion of home range outside the wetland (*HR*0*W*).

We defined for each individual-year a wetland of interest (*WI*) that has been compared with each individual-year home range to construct our metrics. We began by merging the areas modelled by the four methods for each individual-year and then the total wetland was intercepted by the merger (**Fig. 2. a**). For each method (for example the Kernel method in **Fig. 2. b.c**) we then calculated the *HR0W* (expressed in percentage, %), obtained by dividing the surface of the home range outside the *WI* contours by the total surface of the home range modelled by each method (**Fig. 2. c**).

**Fig. 2:**
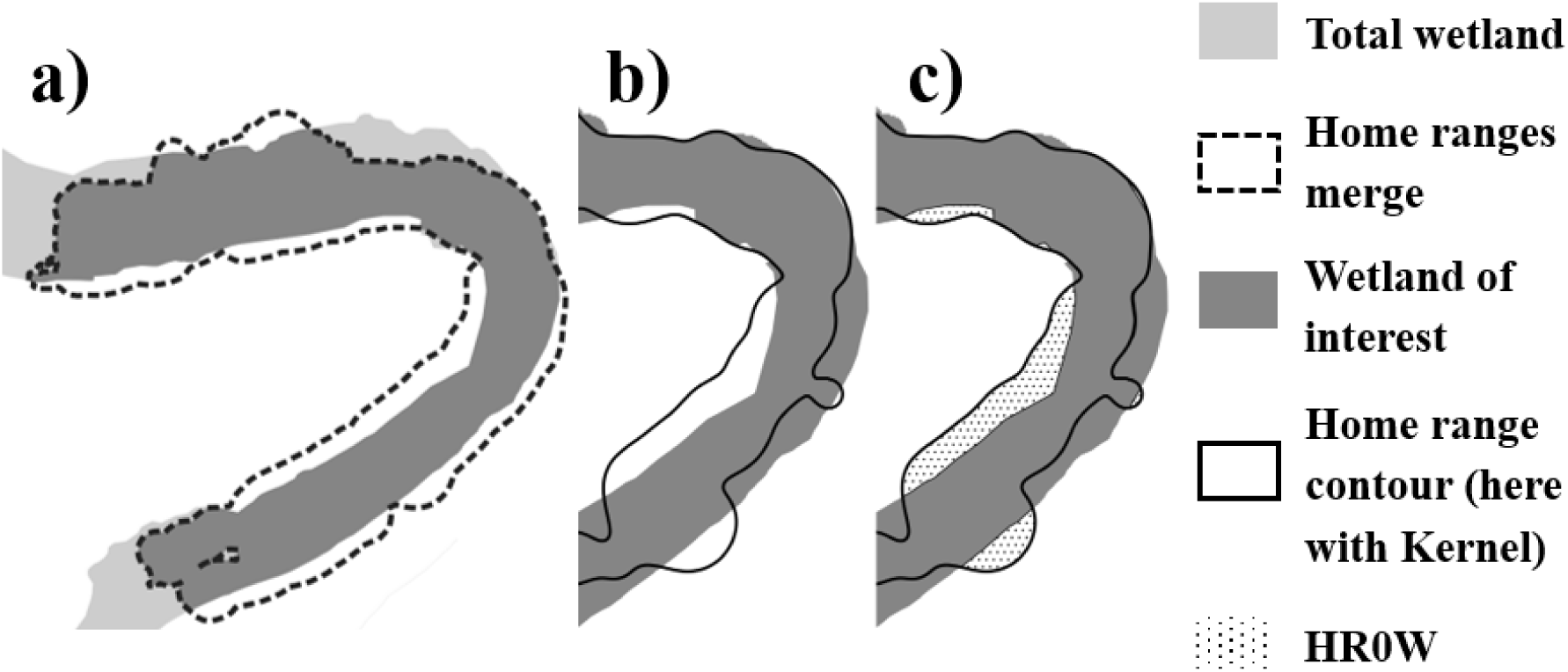
Examples of modelling the different surfaces used to measure HR0W (Proportion of home range outside the wetland). a) modelling of the WI (Wetland of interest) where light grey area is the total wetland, dotted line represents the contours of the merge of the four methods home ranges, and dark grey is the WI; b) comparison of the Kernel home range with the WI to find c) the HR0W.

We selected as the “best” method the one with the lowest HR0W, depending on the landscape.

### 2.6. Data analysis

We excluded individual-years tracked less than one month (3♂ in LG, 2♀ in CA). We discarded three individuals with < 65 relocations and focused on the 16-remaining individual-years (threshold explained in ESM 1). All analyses were computed using R (v. 4.4.1, R Development Core team, www.r-project.org), with significance set at p-value < 0.05.

All analyses were computed using R (v. 4.4.1, R Development Core team, www.r-project.org), with significance set at p-value < 0.05.

A Generalized Linear Mixed Model (GLMM) with beta distribution was first fitted to explain *HR0W*, using the glmmTMB package (Brook et al., 2025, v.1.1.11 for R software). Covariates in the starting umbrella model were home range estimation method (4: EHR, GAM, Kernel, a-LoCoH), the landscape type (2: river and marsh) and their two-way interaction. We included individual-years as a random effect to account for repeated measures (e.g each repeated four times, once per method).

To assess variability of home range and core area size after the method was chosen (see above), we used a Linear Mixed-effect Model (LMM) with following covariates: sex, ecological period (breeding or non-breeding), landscape type (if the selected method was consistent across landscapes), sex –period and sex –landscape two-ways interactions if applicable. We defined breeding period as April 1^st^ - August 31^st^, non-breeding as the rest of the year. We associated each individual-year with the period having the most locations. We considered individual as a random effect to account for dependence when monitored over two years.

Linear mixed-effects models were fitted using the lme4 (Bates et al., 2024, v.1.1-35.5 for R software) and the lmerTest packages (Kuznetsova et al., 2020, v.3.1-3 for R software).

From all previous models, we selected the most plausible using the corrected Aikaike Information Criterion (*AICc*) using the *dredge* function of the MuMIn package (Barton et al., 2024, v.1.48.4 for R software). We selected models with *ΔAIC_C_* < 2 as most plausible. If several models present *ΔAIC_C_* < 2, we conducted model averaging with the model.avg() function of the MuMIn package and we used conditional average betas to define the final model. If no betas were significant, we selected the model with the lowest *ΔAIC_C_*. *AIC* tables are presented in ESM 2.

## 3. Results

We used 16 individual-years in analyses (**Table *2***), tracked during a mean of 153 days (min: 50, max: 230) and resulting in a mean of 129 daily locations (min: 48, max: 213), to model home range with the four different methods (**Fig. 3**).

**Figure 3:**
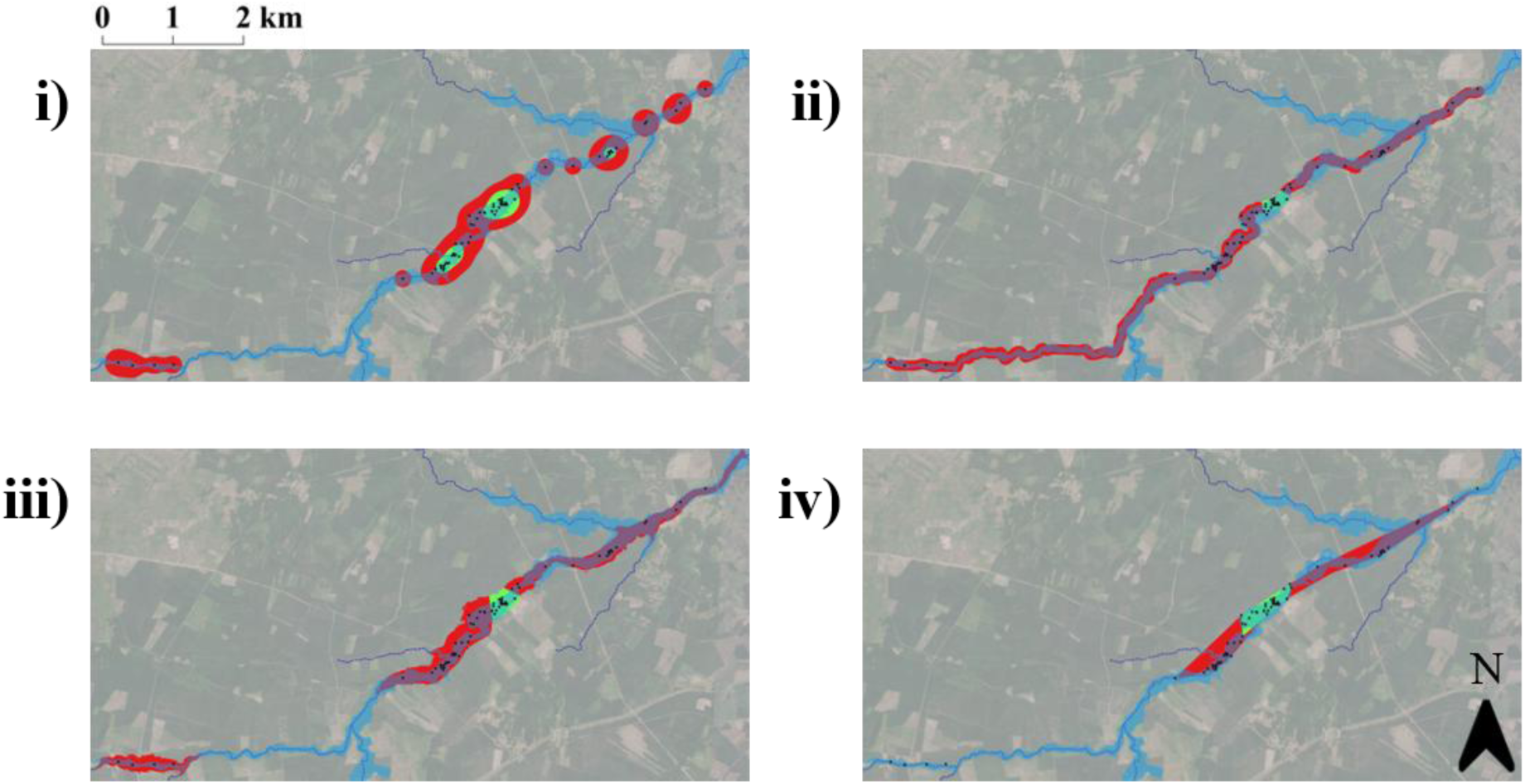
Examples of how the four methods model the home range for the M2C European mink female in a river landscape type. Dark blue dashed lines: watercourse; black points: diurnal locations; light blue shaded area: wetland; red: 95% home range estimated; green : 50% home range estimated with i) Kernel method; ii) EHR method; iii) GAM method; iv) a-LoCoH method.

**Table 2:** Radiotracking data of European mink *Mustela lutreola* in France, used in statistical analyses to estimate home range size.

| Animal ID | Sex | Region | Landscape type | Radiotracking period | Radiotracking duration (days) | Number of diurnal locations | Reason for stopping radio-tracking |
| --- | --- | --- | --- | --- | --- | --- | --- |
| M2C | F | LG | River | 1997/07/24-1998/04/26 | 276 | 122 | End of transmitter life |
| M2C2 | F | LG | River | 1998/12/11-1999/03/19 | 98 | 80 | End of transmitter life |
| M3C | F | LG | River | 1997/12/19-1998/04/12 | 114 | 82 | End of transmitter life |
| M4C | F | LG | River | 1998/01/06-1998/03/26 | 79 | 73 | End of transmitter life |
| M5C | M | LG | River | 1998/03/14-1998/10/08 | 208 | 162 | End of transmitter life |
| M10C | M | LG | River | 1999/03/06-1999/08/06 | 153 | 107 | End of transmitter life |
| M3E | M | LG | River | 1996/12/26-1997/03/25 | 89 | 77 | End of transmitter life |
| Badu | M | CA | River | 2020/02/16-2020/09/07 | 204 | 163 | End of transmitter life |
| Badu2 | M | CA | River | 2021/02/27-2021/04/18 | 50 | 48 | Predation by fox |
| Gwenn | M | CA | River | 2020/10/18-2021/03/12 | 145 | 139 | Predation by fox |
| Maya | F | CA | River | 2021/03/23-2021/11/08 | 230 | 213 | End of transmitter life |
| Mellea | F | CA | River | 2020/10/25-2021/05/30 | 217 | 211 | End of transmitter life |
| Popeye | M | RM | Marsh | 2020/01/25-2020/06/17 | 144 | 98 | End of transmitter life |
| Tim | M | RM | Marsh | 2020/10/30-2021/05/07 | 189 | 176 | End of transmitter life |
| Naia | F | RM | Marsh | 2020/11/28-2021/06/01 | 185 | 185 | End of transmitter life |
| Hagrid | M | RM | Marsh | 2021/03/04-2021/07/24 | 142 | 133 | End of transmitter life |

### 3.1. Home range proportion outside wetland (HR0W)

Model selection supported an additive model including both estimation method and landscape type as explanatory variables for *HR0W* (ESM 2, Table S3).

The mean *HR0W* (**Fig. 4**) is significantly higher with the Kernel method (mean = 0.38, 95% CI: 0.24-0.52) than with EHR (mean = 0.26, 95% CI: 0.17-0.36, *p* = 0.002) or GAM (mean = 0.21, 95% CI: 0.12-0.31, *p* < 0.001). With the a-LoCoH (mean = 0.34, 95% CI: 0.19-0.49), the mean *HR0W* is higher only compared to the GAM (*p* = 0.01). There are no other significant differences. In addition, *HR0W* is significantly higher in river landscape (mean = 0.39, 95% CI: 0.32 -0.45) than in marsh landscape (mean = 0.02, 95% CI: 0.02-0.04, *p* < 0.001).

**Fig. 4:**
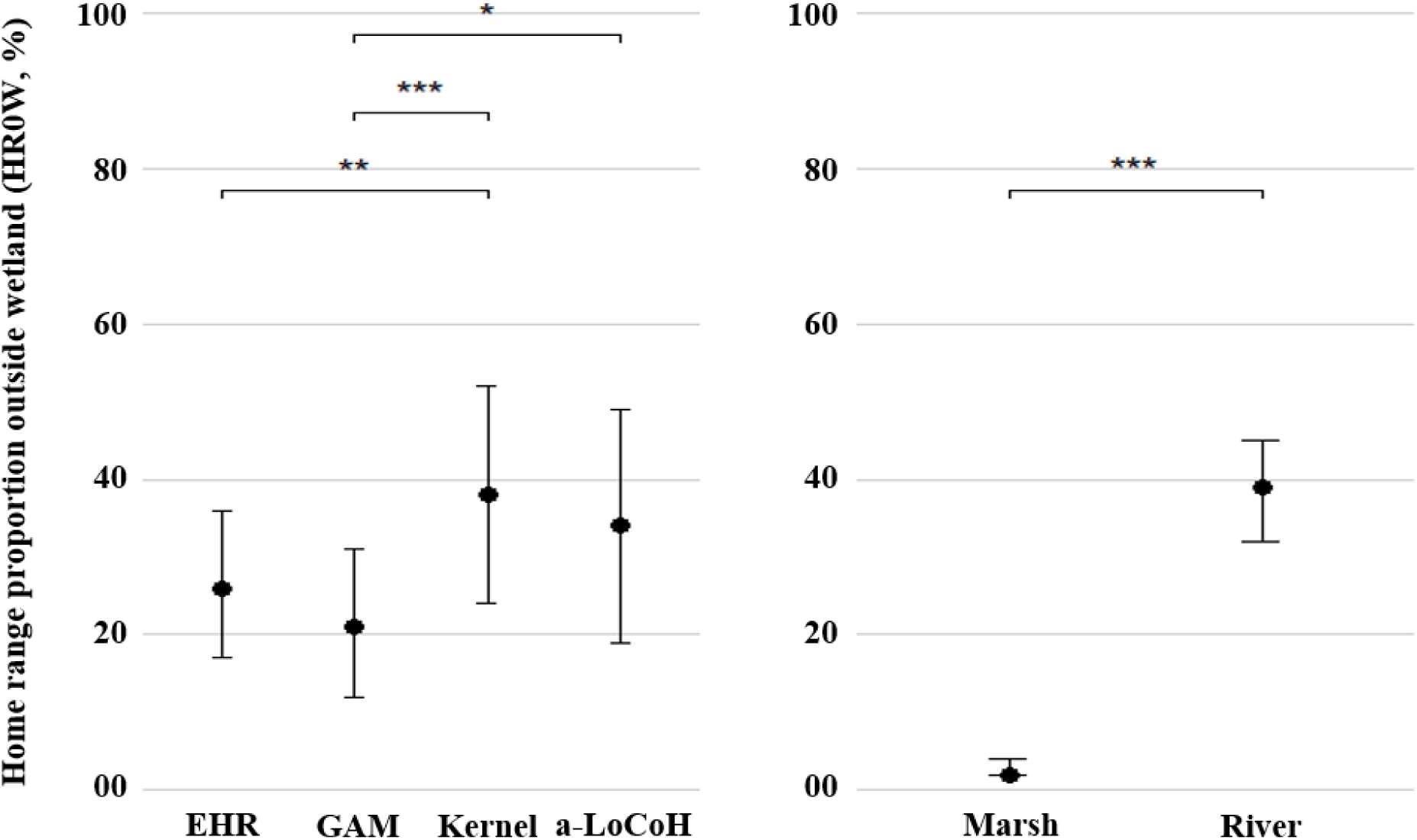
Observed home range proportion outside wetland (*HR0W*) for the four home ranges modelling (left) methods (EHR: Ecological Home Range; GAM: Generalized Additive Model Kernel, a-LoCoH) and for the two landscapes type (right). Filled dots: mean value for each method or landscape; bars: standard deviation. *p ≤ 0.05; **p ≤ 0.001; ***p ≤ 0.0001.

As a synthesis, we reject the Kernel method for European mink home range estimation, due to significant higher *HR0W* than GAM and EHR. We also reject a-LoCoH because higher HR0W than GAM. EHR and GAM are similar in terms of HR0W. As EHR is a method that cannot be improved whereas GAM method offers the suitability to add variables to precise ecological requirements (see discussion below), GAM method is used preferentially below to analyse the pattern of variability of the home range size.

### 3.2. Home range and core area size using the GAM method

Three models including the simple effects of sex, the addition of landscape type and sex as well as null model were equally supported by the *AIC_C_* model selection (ESM 2, Table S5). Model averaging estimates (conditional average) showed that landscape type and sex was significant, so the chosen model was the addition of landscape type and sex as explanatory variables. Home range size is significantly larger for males (mean = 3074.18 ha, 95% CI: 1186.63 ha-5308.47 ha; range: 298.0 ha –8760.9 ha) than for females (mean = 116.45 ha, 95% CI: 60.30 ha-175.38 ha; *p* = 0.01; range: 28.4ha –256.9) and is significantly larger in river landscape (mean = 2251.02 ha, 95% CI: 776.20 ha-4219.62 ha) than in marsh landscape (mean = 367.63 ha, 95% CI: 174.10 ha-561.16 ha, p = 0.048) (**Fig. 5**).

**Fig. 5:**
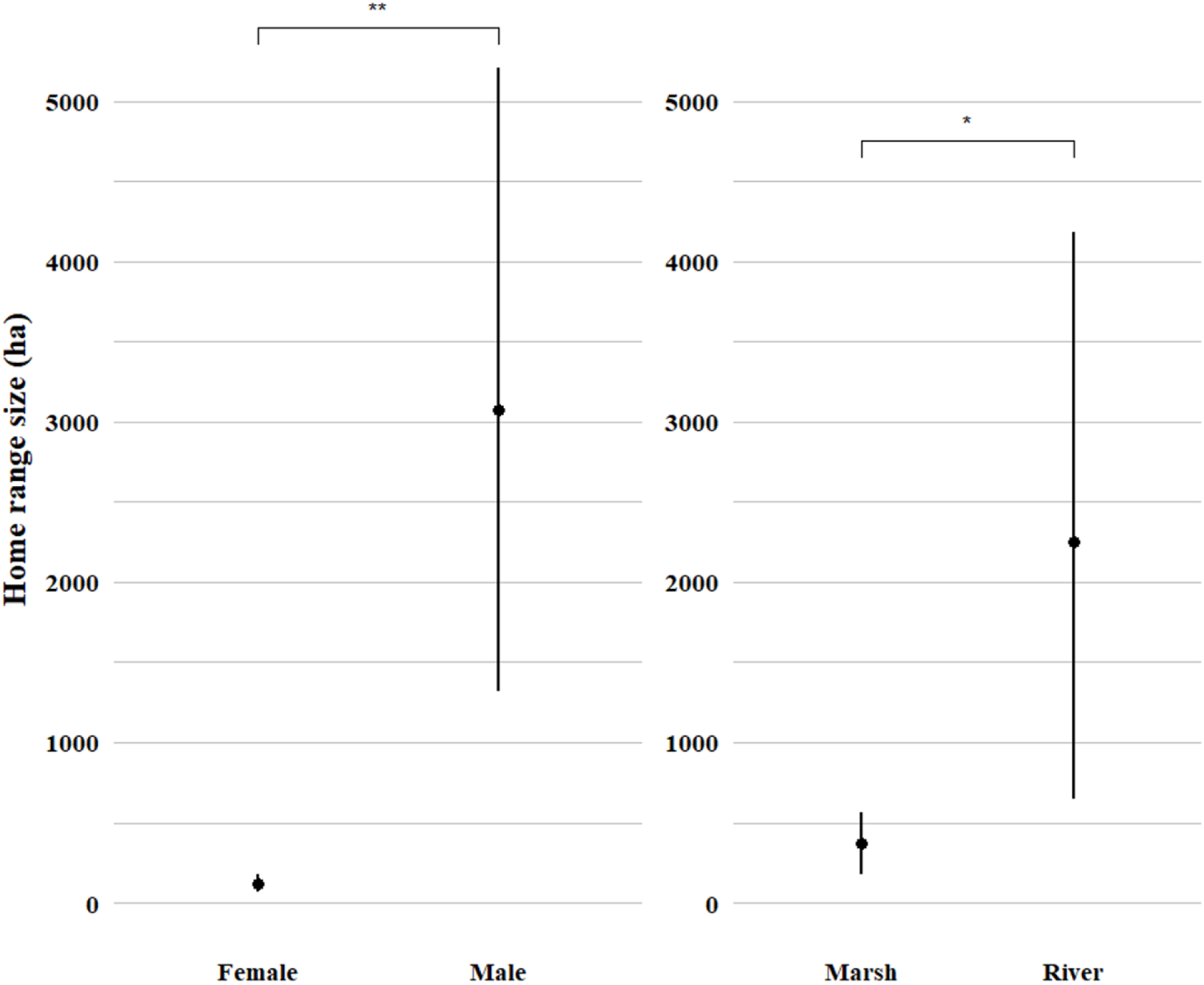
Home range size using the GAM method between sex (left) and landscapes type (right). Dots: mean value for each method or landscape; bars: standard deviation. *p ≤ 0.05; **p ≤ 0.01; ***p ≤ 0.001.

Concerning the variability of the core area size, the best model was the null model, and two models with the sex and the period as independent variables were equally supported (ESM 2, Table S6). However, model averaging shows that none of these two variables is significant, suggesting that the core area size is not explained by any of our predictors. Values of home range and core area size using the GAM method are presented in **Table 3**.

**Table 3:** Home range and core area size for the 16 individual-years using the GAM method.

| Animal ID | Sex | Region | Landscape type | Number of diurnal locations | Home range size (ha) | Core area size (ha) |
| --- | --- | --- | --- | --- | --- | --- |
| M2C | F | LG | River | 122 | 161.7 | 12.0 |
| M2C2 | F | LG | River | 80 | 129.6 | 24.2 |
| M3C | F | LG | River | 82 | 28.4 | 1.5 |
| M4C | F | LG | River | 73 | 51.1 | 4.6 |
| M5C | M | LG | River | 162 | 8760.9 | 1816.4 |
| M10C | M | LG | River | 107 | 2622.6 | 250.4 |
| M3E | M | LG | River | 77 | 1214.2 | 142.3 |
| Badu | M | CA | River | 163 | 2862.4 | 56.4 |
| Badu2 | M | CA | River | 48 | 2808.7 | 174.8 |
| Gwenn | M | CA | River | 139 | 7978.4 | 457.2 |
| Maya | F | CA | River | 213 | 256.9 | 46.2 |
| Mellea | F | CA | River | 211 | 137.2 | 45.0 |
| Hagrid | M | RM | Marsh | 133 | 571.0 | 124.1 |
| Naia | F | RM | Marsh | 185 | 50.2 | 8.2 |
| Popeye | M | RM | Marsh | 98 | 551.3 | 121.3 |
| Tim | M | RM | Marsh | 176 | 298.0 | 125.1 |

## 4. Discussion

In order to identify a relevant method for home range size estimation accounting for spatial specialisation and daily locations of semi-aquatic mammals, whose movements are constrained by the dendritic systems in which they live, we compared the home ranges of 16 European minks radiotracked in France in two landscapes types (river and marsh), modelled by two novel methods for such species (Ecological Home Range, EHR, and Generalized Additive Model, GAM) with a highly-used control, the Kernel method, and a well-tested spatial constraint control, a-LoCoH. Using the proportion of home range outside the wetland (*HR*0*W*), our findings revealed that the four methods differed in their adaptation to wetland environments. It was obvious that the Kernel method was not suitable for estimating the home range of the European mink as it did not match well with the wetland. For the a-LoCoH, we set this method aside, given that HR0W was significantly higher than the GAM and not significantly different than the Kernel which had the highest HR0W (meanwhile the worst).. In fact, a-LoCoH was not well-fitted for the European mink, using as **a** the heuristic value given by Getz et al. (2007). Choosing between the EHR and GAM methods as the “gold standard” for European mink home range estimation remained a matter of debate. Indeed, whilst the GAM differs significantly from the Kernel and the LoCoH, the EHR appears similar to the latter according to our metric. Differences appear to be influenced by landscape type, suggesting that European mink may use space differently in river versus marsh landscapes. Sex also influenced home range size, while no seasonal effect was observed.

### 4.1. Most suitable home range estimation method

The Kernel method effectively served as a negative control, consistently producing a high proportion of home range areas extending beyond the wetland or the riparian landscape. In some cases, more than 50% of the estimated home range fell outside suitable areas, especially in meandering rivers. Similar overestimations were reported by Tarjan et al. (2016) on sea otters (*Enhydra lutris*), where Kernel-based home range overlapped with inland areas not used by the species. These findings reflect the method intrinsic limitations: assuming isotropic space use around each location, it generates smoothed contours that may suit homogeneous environments but fail for species constrained to dendritic features. Consequently, Kernel often includes ecologically irrelevant zones as observed by Fournier et al. (2008), Slaght et al. (2013), and Leite et al. (2016). For the critically endangered European mink, such unrealistic representations may mislead conservation planning and hinder the identification of priority areas, including in reintroduction.

Our findings on a-LoCoH did not allow us to distinguish it from Kernel, which is rather quite surprising given that LoCoH, and especially adaptative LoCoH, is known to be more effective than Kernel to not model unused areas (Getz et al., 2007; Lichti and Swihart, 2011). We found that there were no differences in the proportion of home range outside the wetland between a-LoCoH and Kernel. An explanation to these results could be the number of locations used in our study. In fact, we modelled home range with a range of 48 to 213 locations. In their description of the method, Getz et al. (2007) analysed the effect of sample size on modelling error. To this end, they use samples ranging from 200 to 1,000 data points. The lowest value they used is the highest value in our study. In their study, Socia-Martinez et al. (2023) also demonstrated that the LoCoH method requires a large amount of data to be effective. For a species as elusive as the European mink, for which data collection is very difficult, it appears that the LoCoH method may not be suitable.

Determining a “gold standard” between the EHR and GAM methods is more complex, as both appear not significantly different, but significantly lower than the Kernel.

The EHR method aims to provide spatially realistic home ranges of species strictly dependent on wetlands. Here, we refined previous approaches by defining, for each individual, a personalized buffer around the hydrographic network based on their observed locations, rather than a uniform buffer (e.g., Korbelová et al., 2016; Halbrook and Petach, 2018). This flexible approach computes distance between each location and the nearest watercourse, producing continuous home ranges from upstream to downstream. Compared to other constrained methods such as the Adjusted Home Range (AHR; Palomares et al., 2017b), EHR offers a simpler and time-efficient alternative, avoiding intensive spatial features mapping. Its reproducibility makes it especially useful when high-resolution data are unavailable. However, it depends heavily on the accuracy of the hydrographic dataset, offers limited flexibility for parameter tuning, and lacks internal fit metrics. Moreover, including all watercourses between extreme locations may incorporate unused. A possible improvement could be to preselect only watercourses with actual locations.

The GAM approach estimates home ranges from predicted probabilities of presence derived from locations and environmental variables. Unlike geometric or buffer-based methods, it incorporates resource selection into space-use estimation. By modelling presence probability as a function of spatial coordinates and environmental covariates, it yields ecologically informed home range, similar to the synoptic model of Horne et al. (2008) applied to riverine species like Blakiston’s fish owl (Slaght et al., 2013) or the water opossum (Leite et al., 2016). In our study, GAM integrated species distribution modelling with home range estimation. One of the main benefits of such an approach lies in its ability to generate home ranges that reflect the fine-scale spatial and environmental constraints shaping the animals’ movements. In our case, the model effectively integrated spatial position with variables tied to the species’ ecology, producing home ranges that were continuous where suitable but also responsive to spatial structure. Its flexibility allows adding variables such as vegetation structure, land cover, or indicators of anthropogenic pressure. Yet, it requires high-quality environmental data, careful selection, and statistical expertise. Unlike EHR, it may predict disjointed high-probability zones, potentially omitting travel corridors. Despite these caveats, GAM provides a powerful and ecologically grounded method. Among the methods evaluated, we recommend GAM as the most suitable for species constrained to dendritic systems such as wetlands. It should be noted that, in order to apply the GAM and the EHR to other species, prior knowledge of the ecology of that species—and in particular of the variables that constrain its home range—must be obtained.

### 4.2. Home range size

In our study, the average home range size much larger for males than for females. These values represent the largest home ranges ever reported for the European mink. When comparing absolute sizes, male home ranges in our study are comparable to those reported by Fournier et al. (2008) (2,971 ± 1,888 ha, *n* = 3), but significantly larger than those observed by Palomares et al. (2017b) (MCP: 253.2 ± 28.1 ha; AHR: 78.4 ± 7.3 ha, *n* = 21) and Ceña et al. (2003) (83.7 ± 138.9 ha, *n* = 21), being respectively 22, 39 and 36 times larger. For females, the discrepancy is also pronounced and contrasted: their home ranges in our study are approximately twice smaller than those reported by Fournier et al. (257 ± 113 ha, *n* = 4), 3 to 7 times larger than those of Palomares et al. (2017b) (MCP: 40.4 ± 7.4 ha; AHR: 17.4 ± 2.1 ha, *n* = 21), and roughly three times larger than those of Ceña et al. (34.9 ± 43.5 ha, *n* = 25). Moreover, female home ranges were only 4% of male home ranges. This sexual dimorphism in home range size is more pronounced than in previous studies. Fournier et al. (2008) reported that female home ranges were 9% of male home ranges, Palomares et al. (2017b) 16% (AHR) and 22% (MCP), and Ceña et al. (2003) observed the smallest difference, with females’ home ranges being 42% of male ranges.

These huge differences in home range sizes may reflect contrasts in population densities between French and Spanish populations. For instance, Ceña et al. (2003) monitored 167 km of rivers in Spain over three years and captured 48 individuals for 16,861 trap-nights. In comparison, in the four-year study in the Landes de Gascogne, with a similar sampling effort (14,731 trap-nights) but over a smaller area (68 km of rivers), resulted in only 11 captured individuals. A similar trend is observed when comparing studies in Navarre (Palomares et al., 2017b) and Charente (our data). In Navarre, Palomares et al. (2017b) monitored 60 km of rivers over three years and captured 28 individuals for 877 trap-nights. Meanwhile, in Charente, the three-year study over 40 km of rivers, with a much greater effort (4,498 trap-nights), resulted in only 5 captured individuals.

These comparative figures strongly suggest substantially lower densities of European mink in France than in Spain, which likely contributes to the larger home range sizes observed in our study. This pattern aligns with established ecological theory that predicts an inverse relationship between population density and home range size in territorial carnivores (Benson et al., 2006; Dahle and Swenson, 2003; Rich et al., 2012).

Alternatively, but not mutually exclusive, it is also possible that the environmental conditions are completely different between the two countries, and that the European mink finds more resources within a smaller area in Spain than in France. The Spanish European minks should exploit smaller landscapes where prey is abundant; this aligns with the habitat productivity hypothesis (Nilsen et al., 2005), which proposed that a rich environment allows animals to meet their metabolic needs over a smaller area. Although comparing habitat suitability and productivity relevant to mink in the two countries is beyond the scope of this study. This remains a hypothesis that would need to be tested, as no current data or references confirm a difference in habitat between the two countries.

Finally, although the studies against which we are comparing home range sizes used estimation methods that took into account the ecological constraint of specialisation in wetland habitats, none of them used the GAM, which may lead to certain differences. Fournier et al. (2008) used the “Cluster analysis” method with the nearest-neighbour rule, developed by Kenward (1987), who defined this method as the most suitable for fitting complicated home ranges from telemetry data. This study by Kenward (1987) subsequently inspired Getz et al. (2007)’s work on developing the LoCoH method. This method therefore potentially has the same drawbacks as those we discussed in section 4.1 on LoCoH, including, amongst other things, the need for a large amount of data in order to be effective. Ceña et al. (2003) used two different methods depending on whether the individuals were found in riverine or marsh areas. In marsh areas, they used a MCP, whilst in riverine areas they defined the home range as the length of the river stretch used (the distance between the upstream and downstream locations) multiplied by the average distance between the locations and the river’s midline. This method is very similar to our EHR, except that we used a proxy for the maximum distance whilst they used the average distance. This could explain the difference in home range size among females (a factor of 3), but not for males (a factor of 36). Finally, Palomares et al. (2017b) initially used a MCP, which has the shortcomings mentioned in the introduction. They also developed a method which they termed AHR (Adjusted Home Range). This method involves delineating the surface occupied by rivers, tributaries, irrigation ditches, lagoons, and their associated riverside vegetation, inside the MCP, using high-resolution ortho-images. Whilst this method must accurately reflect the species’ ecological constraints, it requires a huge amount of time for photointerpretation, particularly if the individuals have a very extensive home range.

### 4.3. Conclusions

Our findings can be applied to the conservation of the species, firstly because the recorded home-range sizes suggest that individuals travel over long distances and are therefore more likely to be exposed to threats such as road collisions and predation. These estimates can also be used to guide reintroduction programmes by informing the number of individuals that can be released within a given area. More broadly, the extensive spatial requirements observed in European mink highlight the need for conservation actions that maintain ecological connectivity, facilitate safe movements, and promote successful dispersal across landscapes. Furthermore, although the space utilised –and therefore the home range –of the European mink is constrained by factors linked to water availability, it may be that differences in occupation of some areas within the home range (core areas) are due to the availability of other habitats. Indeed, it may be the availability of areas with dense vegetation (Zabala et al., 2003; Fournier et al., 2007, Palomares et al., 2017a) that enables the species to find resting sites more effectively, which would therefore explain the spatial distribution of the core areas. Therefore, improving our understanding of habitat requirements and habitat selection should be considered a priority for the conservation of the species. These considerations highlight the urgent need for an updated analysis of habitat use and selection by the European mink, employing modern analytical tools. Even if the three Natura 2000 translocation sites were identified as suitable for the European mink in terms of habitats, such a precise study with new analytical tools, for example by adding habitats parameters to the GAM, would be important in order to provide clear guidance on the quality and appropriateness of these areas for long-term conservation success.

## CRediT authorship contribution statement

**Rémi Bodinier:** Writing –review & editing, Writing –original draft, Visualization, Investigation, Formal analysis, Data curation, Conceptualization. **Christine Fournier- Chambrillon**: Writing –review & editing, Validation, Supervision, Methodology, Investigation, Conceptualization. **Stéphane Aulagnier:** Writing –review & editing, Methodology, Investigation. **Yoann Bressan:** Writing –review & editing, Investigation. **Romain Beaubert:** Writing –review & editing, Investigation, Data curation. **Sébastien Devillard:** Writing –review & editing, Validation, Supervision, Project Administration, Funding acquisition, Conceptualization. **Pascal Fournier:** Writing –review & editing, Validation, Supervision, Methodology, Investigation, Project Administration, Funding acquisition, Conceptualization.

## Acknowledgments

We would like to thank all field workers who participated in data collection during the radiotracking campaigns: especially C. Maizeret for its coordination of the first program, S. Cardonne, J.-P. Chusseau, B. Delpart, J. Dupuch, T. Gatelier, A. Gigougnoux, D. Jimenez, J. Joachim, K. Lamarque, D. Lanusse, D. Larrieu, N. Piat, F. Spitz for the first campaign, L. Agat, C. Baduel, V; Barret, T. Berti, T. Beshers, O. Bruneau, J. Cazaillon, M. Dupuy, S. Fagart, C. Fagot, L. Ferrand, N. Froustey, C. Galy-Fajou, E. Isère-Laoué, L. Jomat, V. Mahu, I. Marchand, A. Meunier, B. Ollivier, P. Rigou, C. Wagner, for the second. We also thank Clément Calenge for his expertise on home range estimation methods and for guiding us in our research.We also thank the French Ministry for the Ecological Transition through the National Action Plans concerning European mink, and the Charente, Charente-Maritime, Dordogne, Gironde and Landes departments for their technical help in the field. A preprint version of this article has been peer-reviewed and recommended by PCI Ecology (https://doi.org/10.24072/pci.ecology.101202; Walczyńska, 2026).

## Declaration of Generative AI and AI-assisted technologies in the writing process

During the preparation of this work, the authors used ChatGPT to polish the text, enhance the English language, and reduce the length of an advanced draft to meet the journal’s manuscript length requirements. After using this tool, the authors reviewed and edited the content as needed and take full responsibility for the content of the published article.

## Electronic Supplementary Material 1: Selection method of individuals-years included in analysis

For each individual monitored for more than a month, we randomly selected a number of locations in increments of ten, ranging from ten to the tenth below the total number of locations. For each group of ten locations, we repeated the selection process thirty times. We modelled the Minimum Convex Polygon 100% (Mohr, 1947) home range for each selection and measured its size. The average size of the home range was then calculated for the thirty iterations within each group of ten. We plotted the average home range size as a function of the number of selected locations. We visually assessed whether the plot approached an asymptote and identified individuals for which this was the case. This resulted in the selection of 12 individual-years (**Figure S7**). For these 12 individual-years, we plotted the logarithm of home range size against the logarithm of the number of locations. We calculated the equation of the resulting plot and determined the number of locations needed to model a home range that covered 90% of its total size. Finally, we calculated the average number of locations required across the 12 individual-years and selected all other individual-years with a total number of locations exceeding this mean. It resulted in the selection of four more individual-year for a total of 16 individual-years used for the data analysis (14 different individuals, including two for which two periods over two years of monitoring are available).

**Figure S6:**
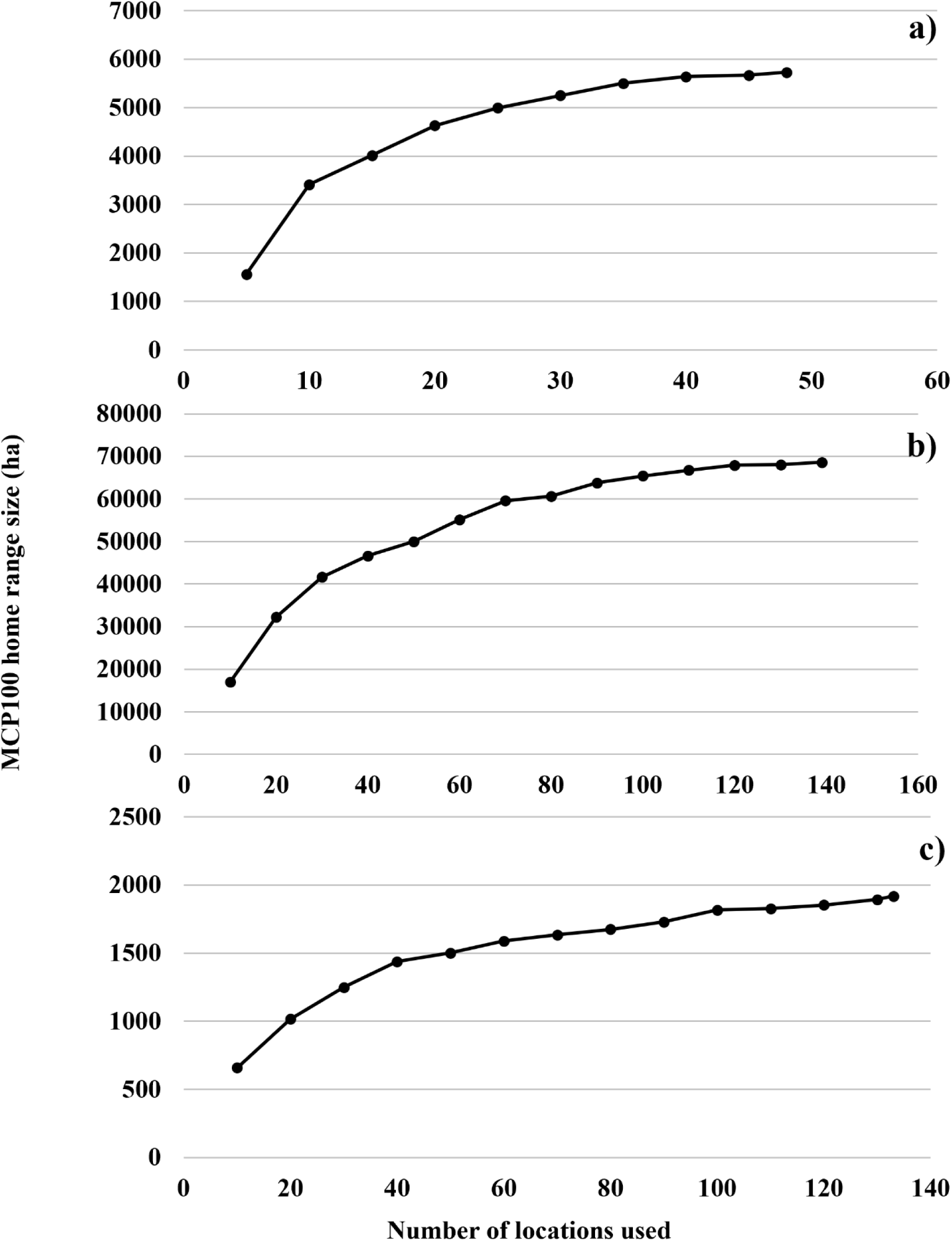
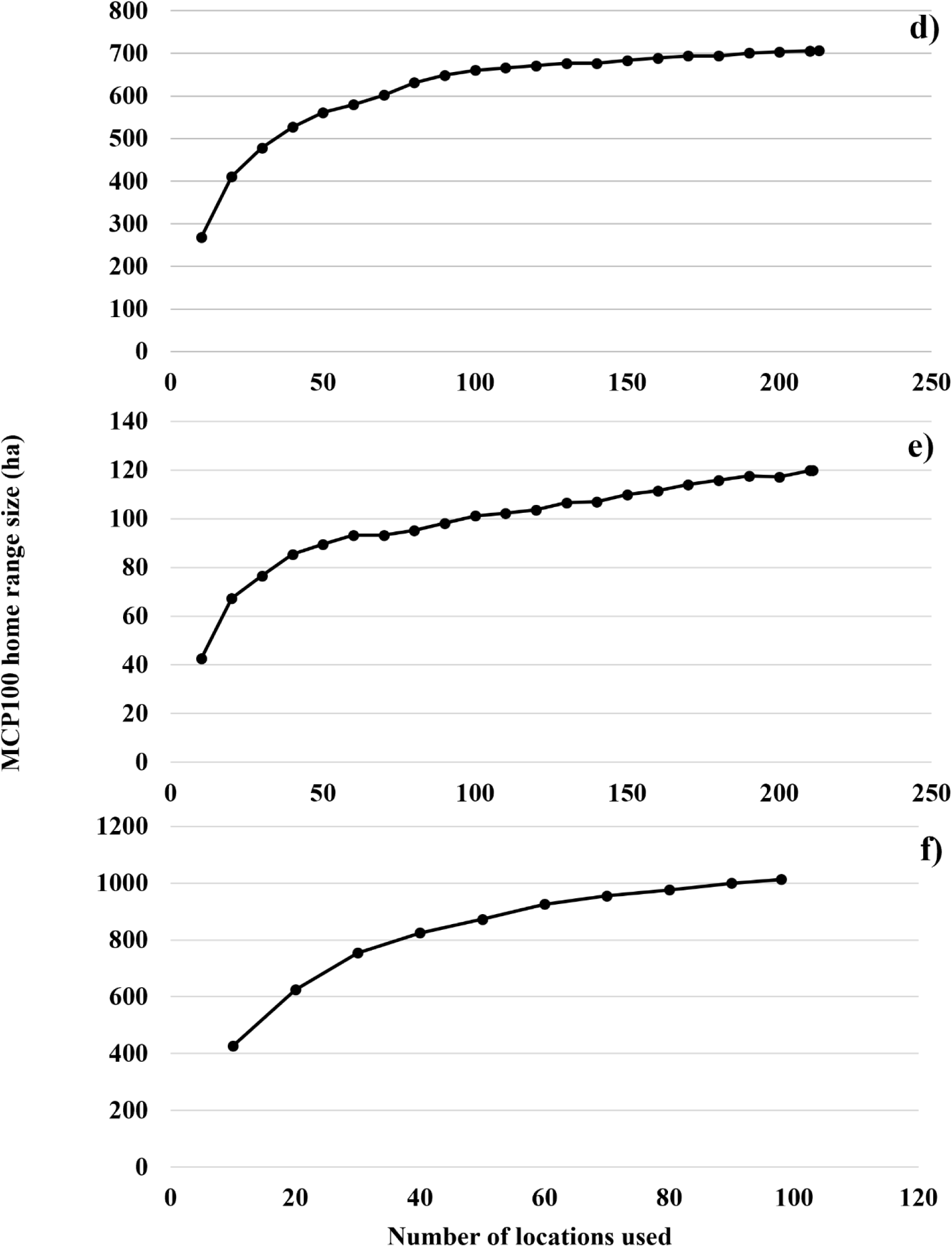
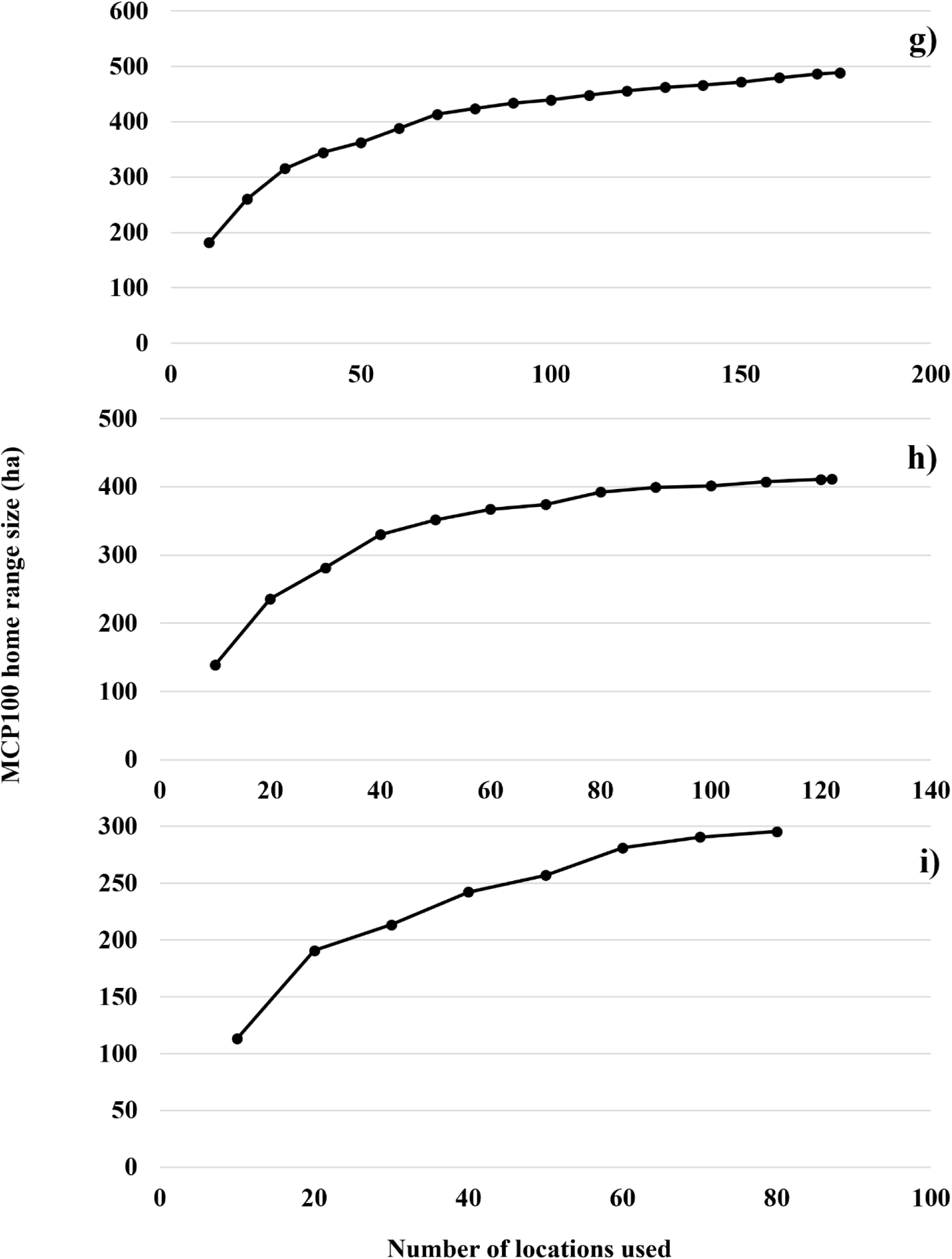
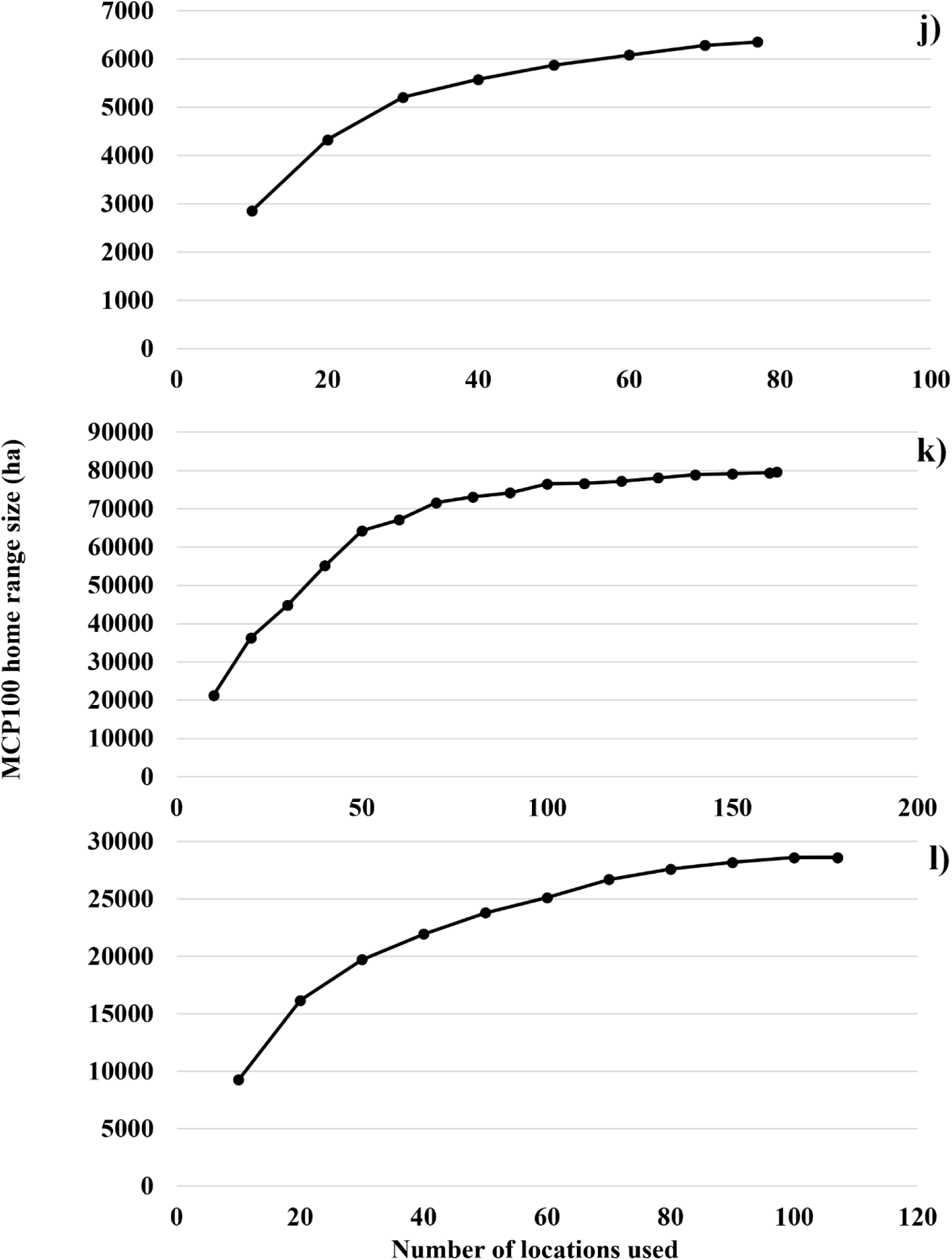
Plots of the average home range size as a function of the number of selected locations for each individual-year that approached visually an asymptote. a) Badu2, b) Gwenn, c) Hagrid, d) Maya, e) Mellea, f) Popeye, g) Tim, h) M2C, i) M2C2, j) M3E, k) M5C, l) M10C.

## Electronic Supplementary Material 2: *AIC* tables for all models used in this article

**Table S4:** Model structure, Akaike’s Information Criterion adjusted for sample size (*AIC_C_*), differences in *AIC_C_* (*ΔAIC_C_*), and Akaike weight (*w_i_*) for the four models predicting the HR0W (in bold, the selected model).

| Variables contained in the model | $AIC_C$ | $\Delta AIC_C$ | $w_i$ |
| --- | --- | --- | --- |
| Landscape type + Estimation method | **-110.3** | **0.00** | **0.902** |
| Landscape type + Estimation method +<br>Landscape type:Estimation method | -105.6 | 4.62 | 0.090 |
| Estimation method | -100.3 | 9.91 | 0.006 |
| Landscape type | -97.7 | 12.57 | 0.002 |
| Null model | -87.7 | 22.58 | 0.000 |

**Table S5:**
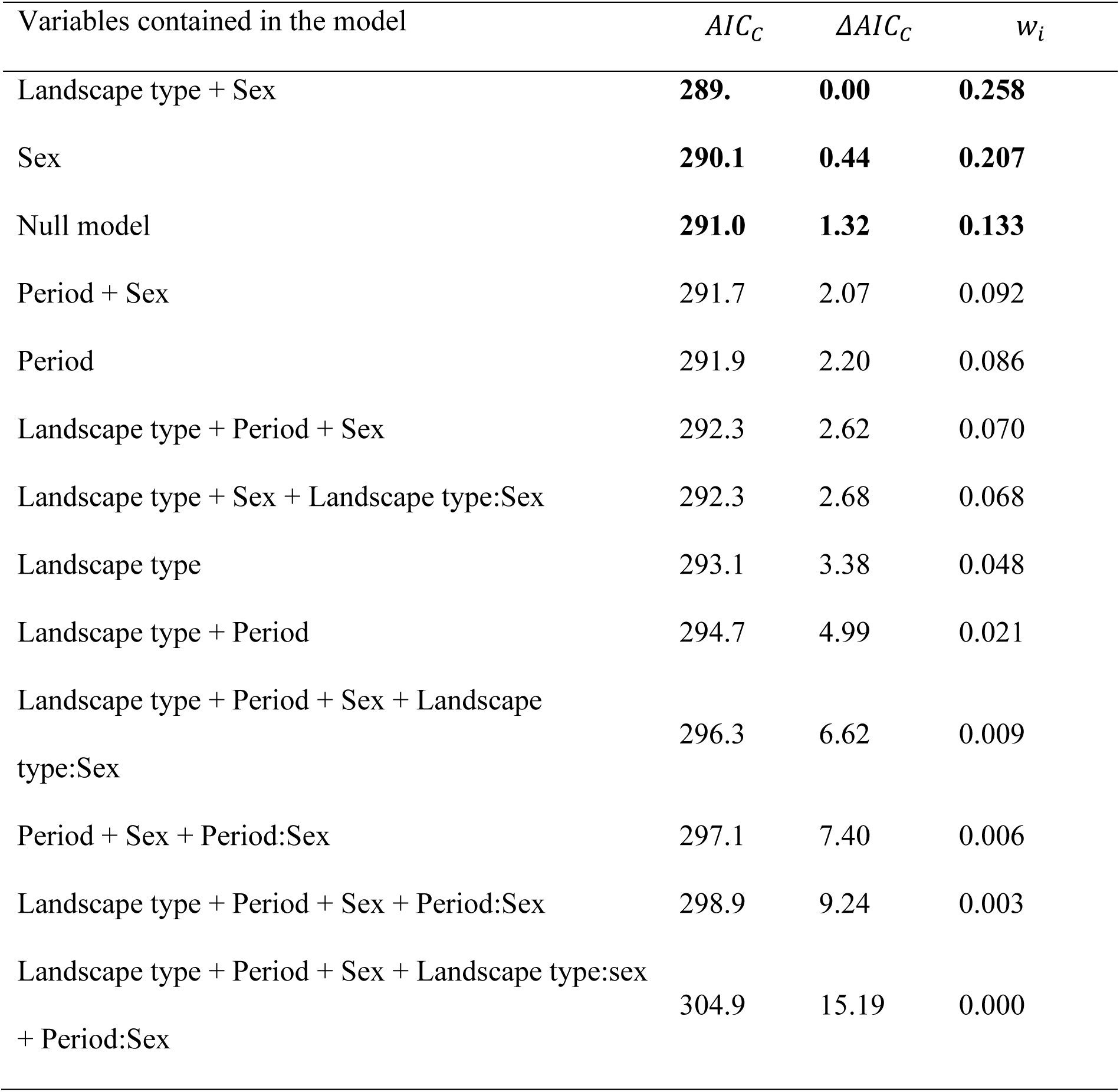
Model structure, Akaike’s Information Criterion adjusted for sample size (*AIC_C_*), differences in *AIC_C_* (*ΔAIC_C_*), and Akaike weight (*w_i_*) for the twelve models predicting the home range size using the GAM method (in bold, the models supported by the selection).

**Table S6:** Model structure, Akaike’s Information Criterion adjusted for sample size (*AIC_C_*), differences in *AIC_C_* (*ΔAIC_C_*), and Akaike weight (*w_i_*) for the twelve models predicting the core area size using the GAM method (in bold, the models supported by the selection).

| Variables contained in the model | $AIC_C$ | $\Delta AIC_C$ | $w_i$ |
| --- | --- | --- | --- |
| Null model | **244.3** | **0.00** | **0.269** |
| Sex | **244.7** | **0.35** | **0.227** |
| Period | **244.8** | **0.50** | **0.210** |
| Period + Sex | 246.8 | 2.51 | 0.077 |
| Landscape type | 247.0 | 2.67 | 0.071 |
| Landscape type + Sex | 247.1 | 2.73 | 0.069 |
| Landscape type + Period | 248.1 | 3.82 | 0.040 |
| Landscape type + Period + Sex | 250.2 | 5.92 | 0.014 |
| Period + Sex + Period:Sex | 250.7 | 6.40 | 0.011 |
| Landscape type + Sex + Landscape type:Sex | 250.9 | 6.59 | 0.010 |
| Landscape type + Period + Sex + Landscape type:Sex | 255.0 | 10.71 | 0.001 |
| Landscape type + Period + Sex + Period:Sex | 255.1 | 10.74 | 0.001 |
| Landscape type + Period + Sex + Landscape type:sex + Period:Sex | 261.3 | 17.00 | 0.000 |

## Electronic Supplementary Material 3: Link to find data and script

https://doi.org/10.5281/zenodo.19551616

